# General Regulatory Factors control the fidelity of transcription by restricting non-coding and ectopic initiation

**DOI:** 10.1101/331793

**Authors:** Drice Challal, Mara Barucco, Slawomir Kubik, Frank Feuerbach, Tito Candelli, Hélène Geoffroy, Chaima Benaksas, David Shore, Domenico Libri

## Abstract

The fidelity of transcription initiation is essential for accurate gene expression, but the determinants of start site selection are not fully understood. Rap1 and other General Regulatory Factors (GRFs) control the expression of many genes in yeast. We show that depletion of these factors induces widespread ectopic transcription initiation within promoters. This generates many novel non-coding RNAs and transcript isoforms with diverse stability, profoundly altering the coding potential of the transcriptome. Ectopic transcription initiation strongly correlates with altered nucleosome positioning. We show that Rap1 sterically constrains nucleosomes as its mere binding to the DNA can be sufficient for restoration normal nucleosome positioning, transcription initiation and gene expression. These results demonstrate an essential role for GRFs in the fidelity of transcription initiation and in the suppression of pervasive transcription, redefining current models of their function. They have general implications for the mechanism of transcription initiation and the control of gene expression.

**HIGHLIGHTS:**

- Rap1, Abf1 and Reb1 control the fidelity of transcription initiation and suppress pervasive transcription
- Widespread ectopic transcription initiation in Rap1-deficient cells induces variegated alterations in gene expression
- Altered nucleosome positioning in GRFs-defective cells correlate with ectopic transcription initiation.
- Rap1 controls nucleosomes positioning and transcription initiation at least partially by a steric hindrance mechanism

## INTRODUCTION

Gene transcription originates in promoter regions where general transcription factors assemble to form the transcription pre-initiation complex (PIC), required for the association of RNA polymerase II (RNAPII) to the DNA. PIC requires favorable conditions for assembly, which are generally determined by the action of activators and co-activators. In yeast, after PIC formation RNAPII scans the downstream DNA for the most favourable site of initiation, which does not require RNA synthesis (Fishburn et al., 2016; Murakami et al., 2015). Selection of the transcription start site (TSS) depends on a loose sequence context and generally occurs within 50-100nt downstream of the PIC (Kuehner and Brow, 2006). The chromatin context crucially defines the properties of gene promoters and affects the propensity of RNAPII to initiate transcription.

Active promoters are characterized by regions that are generally depleted in nucleosomes (Nucleosome Depleted Regions, NDR), which is thought to provide access to DNA-binding proteins required for transcription activation (for recent reviews see: Lai and Pugh, 2017; Lieleg et al., 2015). Promoters that are not constitutively active are generally occupied by nucleosomes that are evicted during or immediately before the activation process. NDRs are also positioned at the 3’ end of genes, where they generally coincide with the promoter of a downstream tandemly-oriented gene, and can promote antisense non-coding transcription (Chereji et al., 2018). Promoter NDRs are bordered by two well-positioned nucleosomes one of which is referred to as the +1 nucleosome because it is the first of the array of genic nucleosome that are also generally well positioned and regularly spaced. The concept of the upstream −1 nucleosome is more ambiguous, because the latter can either be the +1 nucleosome of a divergently transcribed gene or a nucleosome that defines the upstream limit of the NDR. This distinction is not merely semantic, because the bona fide +1 nucleosome is characterized by a different histone composition as it typically contains the H2AZ histone variant instead of H2A. Importantly, the +1 nucleosome is associated with the TSS, which is generally located 12-15 nucleotides downstream of its upstream border in yeast (Hughes et al., 2012; Lee et al., 2007; Tsankov et al., 2010; Whitehouse et al., 2007). Formation of the NDR depends on many factors, the weight of which can vary from case to case (for recent reviews see: Lai and Pugh, 2017; Lieleg et al., 2015). One of these factors is the sequence of the DNA that is wrapped around the histone octamer, which defines the most thermodynamically favorable position for nucleosomes on naked DNA. Many studies have however demonstrated that DNA-histone interactions alone do not determine proper nucleosome positioning and that trans-acting factor are required. Among these, Chromatin Remodelers (CR) and General Regulatory Factors (GRFs) play important roles. Among CR, the ones that are the most influential on the architecture and width of the NDR are the SWI/SNF and RSC complexes, which both use the energy of ATP to displace nucleosomes at promoters (Hartley and Madhani, 2009; Parnell et al., 2008; Ryan et al., 1998; Shivaswamy and Iyer, 2008). Other remodelers, like ISWI and INO80 are believed to be important for positioning the +1 nucleosome and the packing of nucleosomes along transcription units (Krietenstein et al., 2016; Lai and Pugh, 2017). General Regulatory Factors (among which Rap1, Reb1, Abf1 and Tfb1) contain a related DNA-binding domain and are required for the expression of several classes of genes, encoding for instance ribosomal proteins, glycolytic enzymes and snoRNAs. GRFs bind the DNA at specific sites generally NDRs and have been shown to be important for excluding nucleosomes from these regions (Badis et al., 2008; Ganapathi et al., 2011; Hartley and Madhani, 2009; Hughes et al., 2012; Kubik et al., 2015; Preti et al., 2010). These proteins do not possess ATPase activity, and therefore they must act on nucleosomes by some different mechanism, either directly or indirectly, for instance by recruiting CRs.

Rap1 (Repressor Activator Protein 1) has been originally described as an activator and a repressor of gene expression (for a recent review see: Azad and Tomar, 2016). Gene activation has been shown to depend on a domain in the C-terminal portion of the protein, which can activate transcription alone when fused to a DNA binding domain with altered specificity (Johnson and Weil, 2017). This region participates to the interaction of Rap1 with PIC components, TFIID and TFIIA (Garbett et al., 2007; Johnson and Weil, 2017; Papai et al., 2010), which has been proposed to be required for transcription activation. Repression of gene expression has been observed at mating type genes and other loci, and it has been suggested that Rap1 negatively affects gene expression by directly interacting with and inhibiting TBP binding to the DNA (Bendjennat and Weil, 2008; Giesman et al., 1991; Kurtz and Shore, 1991; Yarragudi et al., 2004). However, the mechanisms underlying the bimodal role of Rap1 in the control of gene expression are still not fully understood.

One of the salient features of promoters is their intrinsic bi-directionality (Jin et al., 2017; Neil et al., 2009; Rhee and Pugh, 2012; Xu et al., 2009). Besides divergent genes, where bi-directionality is under positive selective pressure, transcription can generally start in both directions, often generating non-coding and non-functional transcription events in the opposite direction of a functional gene (divergent or antisense transcripts). These non-coding transcription events usually generate RNAs that are unstable in wild type cells, which explains why they have remained unnoticed until the advent of genome-wide techniques for analyzing transcription independently of steady state RNA levels. Two main pathways degrade these RNAs with uncharacterized protein coding potential: the first one is nuclear and involves the RNA exosome with its 3’-5’ exonucleases Dis3 and Rrp6; the second one is cytoplasmic and follows the recognition of inappropriately positioned translation stop codons by the nonsense mediated decay (NMD) factors, among which Upf1 (for a review see: Porrua and Libri, 2015).

In spite of the intrinsic bi-directionality of promoters, cells have evolved strategies to direct transcription towards functional coding regions and limit the extent of what is called pervasive transcription (Jin et al., 2017). The nucleosomal architecture of NDRs appears to be important for this control as mutants in chromatin remodelers or modifiers have been shown to increase the extent of pervasive antisense transcription events (Marquardt et al., 2014; Whitehouse et al., 2007). Limiting pervasive transcription is essential for the cell, and spurious transcription events that have escaped control at the level of initiation are terminated by several mechanisms (Porrua and Libri, 2015). Deregulation of either initiation or termination is susceptible to generate interferences that perturb both appropriate gene expression and other DNA-associated events.

In this study we have used a combination of high-resolution genome-wide analyses to address the effects of rapid depletion of Rap1 and other GRFs. We studied the changes in occupancy of RNAPII after rapid Rap1 depletion and correlated these results to changes in transcription initiation, stability of the transcripts vis-à-vis nuclear and NMD degradation pathways, and changes in NDR nucleosome architecture. We demonstrate a massive change in the pattern of transcription initiation, which generates transcripts of diverse stability and coding potential. This translates into variegated effects on gene expression, from activation to repression, and leads to the generation of different protein isoforms, drastically changing current models of GRFs action. We show that Rap1, Abf1 and Reb1 have crucial roles in limiting pervasive transcription at the level of initiation since many novel non-coding RNAs are generated when any of these factors is depleted. Ectopic transcription initiation closely mirrors changes in the distribution of nucleosomes in and around NDRs. We provide evidence that these changes are not due to altered recruitment or function of CR factors by assessing the level and distribution of novel TSSs in CR depleted strains at Rap1 targets. Finally, we demonstrate that the DNA binding domain of Rap1 is sufficient for robust restoration of normal nucleosome positioning, TSS usage and gene expression at many sites. Together, these results support a model whereby Rap1 participates in orchestrating the appropriate pattern of transcription initiation by controlling the position of neighboring nucleosomes, but also by actively preventing spurious initiation within the NDR. This provides a unified view of how Rap1 controls both gene activation and repression, and the quality of the transcripts produced in terms of coding potential. Because we show that neither RNA polymerase occupancy nor overall transcript levels can be considered a priori as faithful predictors of gene expression, our data have important and general implications for the modelling of transcriptional networks.

## RESULTS

### Rap1 depletion promotes ectopic transcription initiation events

In the course of a previous study aimed at describing the function of Rap1 in transcription termination (Candelli et al., 2018) we generated high resolution transcription maps using a modified version of the CRAC (Crosslinking analysis of cDNA) technique (Candelli et al., 2018; Granneman et al., 2009). Briefly, transcription elongation complexes are purified under stringent conditions after UV crosslinking of RNAPII to the nascent RNA *in vivo*. Sequencing of the latter provides high resolution and strand specific information about the position of the polymerase at the moment of crosslinking, bypassing any biases due to the use of RNA abundance as a proxy for transcriptional activity. We first generated CRAC RNAPII transcription maps under conditions of transient depletion of Rap1 from the nucleus with the Anchor Away technique (Candelli et al., 2018; Haruki et al., 2008).

Upon nuclear depletion of Rap1 for two hours, we observed the expected effects of up- and downregulation of RNAPII occupancy at distinct sets of genes containing a Rap1 binding site in their promoter region (Fig. 1A, right of TSS), consistent with the known role of Rap1 as an activator and a repressor of gene expression. A closer look revealed that the depletion of Rap1 is associated with a major global increase in the RNAPII signal in the promoter regions of Rap1 targets, all classes (upregulated, downregulated and unaffected) confounded (Figure 1A, left of TSS border). This increase was observed for at least half of all Rap1 targets as revealed by sorting of the same set of genes based on the CRAC signals upstream of the TSS (Figure 1B).

**Figure 1:**
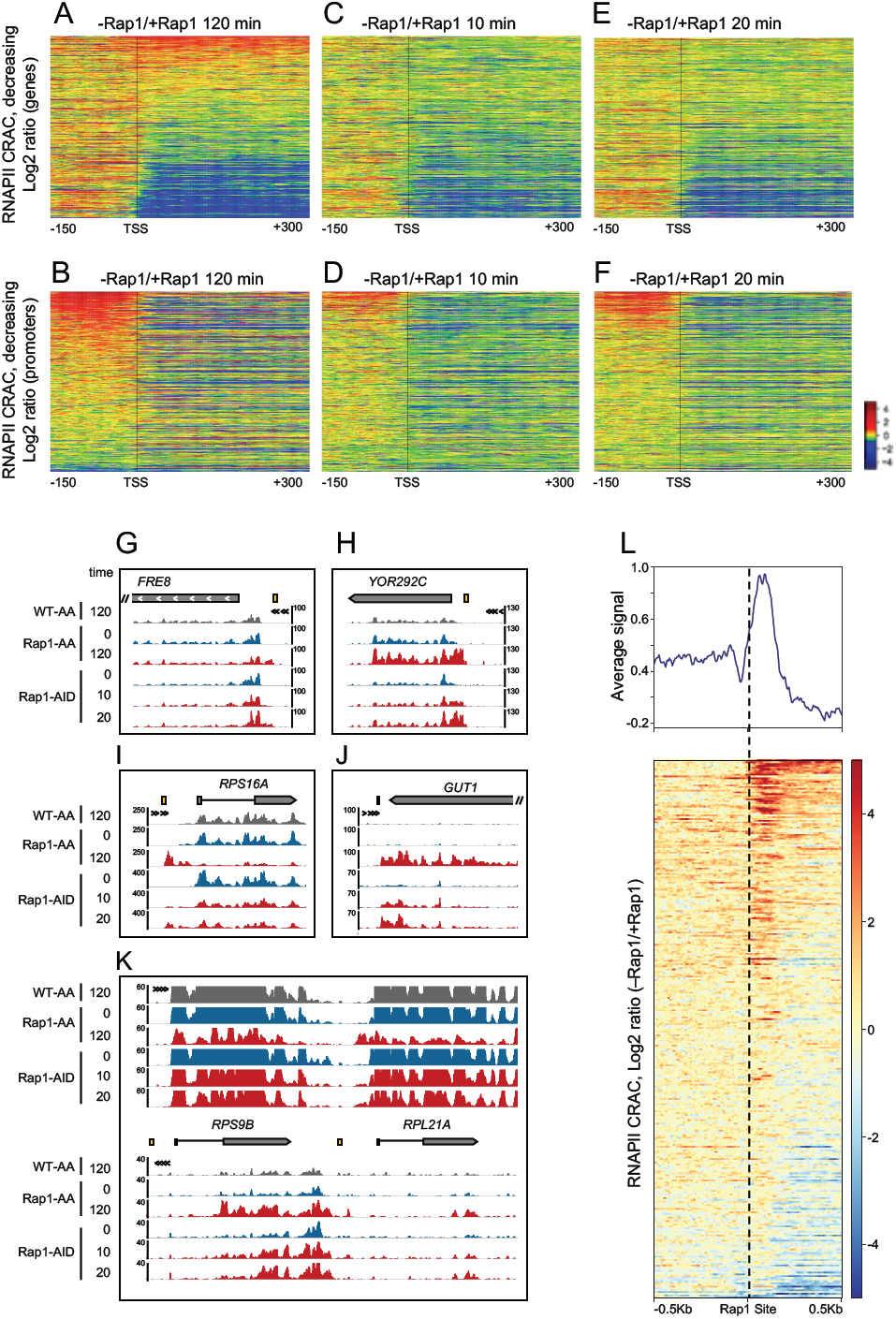
RNAPII CRAC analysis reveals early appearance of ectopic transcription upon Rap1 depletion. See also Figure S1. A-F: heatmaps illustrating the distribution of the Log2 ratio CRAC signal change at 427 Rap1 target genes upon Rap1 depletion by the Anchor-away (A, B) or Auxin-induced degradation (C-F) for the times indicated. Genes have been selected for containing a Rap1 site within 300nt upstream of the TSS. Features are sorted based on decreasing Log2 ratio value in the first 300nt of the gene (A-C) or the promoter region (−150nt upstream of the TSS, D-E). The color codes and the overall signal distribution are indicated. G-K. Snapshots of individual loci illustrating the changes in RNAPII CRAC signal upon Rap1 depletion (red tracks) for the indicated times. Signals derived from Rap1-tagged strains without the addition of the depleting drug or from a non-tagged strain with the addition of rapamycin for 120 min are indicated in blue and grey respectively. The direction of transcription is indicated by multiple arrowheads. The position of the Rap1 site is indicated by a small yellow rectangle. L. Heatmap of CRAC RNAPII signals aligned on Rap1 sites that do not have an mRNA-coding gene within the downstream 500nt. Genomic regions are sorted based on the extent of RNAPII CRAC signal change upon nuclear depletion of Rap1. A metanalysis reporting the average of the signals is shown on top.

We considered that some of the observed alterations in transcription could be due to indirect effects or the co-sequestration of Rap1-interacting factors in the cytoplasm. Therefore we generated genome-wide RNAPII distribution maps after 10 and 20 minutes of Rap1 depletion using the auxin degron system, which allows rapid degradation of a protein of interest (Nishimura et al., 2009a).

Comparison of RNAPII differential profiles sorted as in Figure 1A revealed similar effects on gene downregulation even at very early time points of Rap1 depletion (Figures 1C-E, compare with Figure 1A). With a few notable exceptions that will be described below, genes upregulated after two hours of Rap1 depletion where generally not affected at earlier time points (Figures 1A, C, E and data not shown), suggesting the existence of a late Rap1-dependent direct or indirect phenotype. Importantly, however, increased RNAPII signals in the 150 nt region upstream of gene canonical TSSs appeared as early as 10 minutes of Rap1 depletion and progressively increased at the later time points (Figures 1B, D, F). This was generally concomitant with decreased RNAPII occupancy in the gene body, pointing to a major correlation between these events (see below). Similar clustering of non-Rap1 targets did not reveal a similar increase in RNAPII distribution in promoters, indicating that this effect is specific (Figure S1A-F).

A closer look at individual genes (Figure 1 G-K) revealed that depletion of Rap1 leads to the appearance of RNAPII signals within the NDR, reflecting the unexpected emergence of novel, ectopic transcription start sites (eTSS) at a large (between 30 and 50%) fraction of Rap1 targets. Ectopic initiation was found to arise in one (Figure 1G-J) or both (Figure 1K) directions of transcription relative to the site of Rap1 binding and in a manner that is not dependent on the orientation of the non-palindromic Rap1 site (data not shown). When occurring upstream of canonical transcription units, the presence of eTSS was most often associated to decreased RNAPII CRAC signal in the body of the downstream gene (Figure 1I, Figure S1G-H see also Figure 3H). In some instances, apparent upregulation of transcriptional activity was also observed (Figure 1H and Figure S1I-J; see below), prefiguring diverse effects of Rap1 depletion on gene expression.

**Figure 2:**
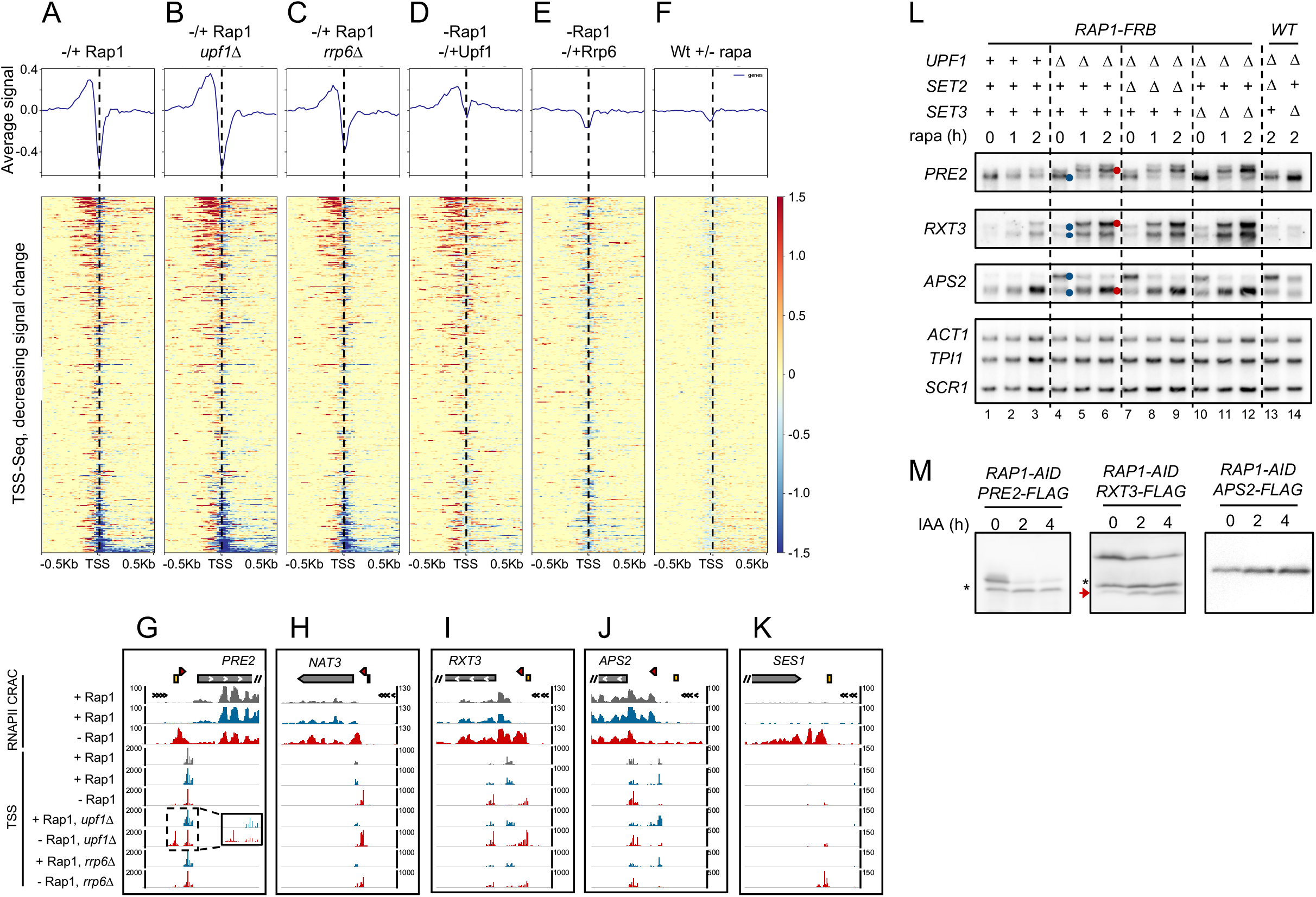
Ectopic transcription initiation impacts gene expression. See also Figure S2. A-F. Heatmaps showing the TSS-seq signal change (Log_2_ratio) upon nuclear depletion of Rap1 (A-C), in the presence/absence of Upf1 (i.e. Rap1-depleted vs Rap1-depleted *upf1*Δ, D) or in the presence/absence of Rrp6 (E). D and E are shown for estimating the extent of unstable RNAs that are produced in Rap1-deficient cells. In F, the same genes are plotted for a control on the impact of rapamycin addition to a non-tagged Rap1 strain. Features are sorted for decreasing change in TSS signals and aligned on the canonical TSS in the wild type strain. G-K Snapshots illustrating the alterations in TSS usage upon depletion of Rap1, with the RNAPII CRAC signal shown in parallel. Track colors for RNAPII CRAC and TSS-seq as in Figure 1. The presence of a red arrowhead in the scheme indicates the existence of a small upstream ORF, which induces NMD sensitivity. L. Northern blot analysis of transcripts produced at the indicated loci upon Rap1 depletion, in different genetic backgrounds. Transcripts produced in the presence of Rap1 are indicated by a blue dot in the *upf1*Δ series of samples. Transcripts that are specifically produced or upregulated in the absence or Rap1 are indicated by a red dot. M. Western blot analysis of proteins produced from HA-tagged *PRE2, RXT3* and *APS2* loci during Rap1 depletion for the indicated times. A red arrow indicates the position of the N-ter truncated isoform of Rxt3 produced when Rap1 is depleted. The asterisk indicates a cross reacting protein that can be used as a loading control.

**Figure 3:**
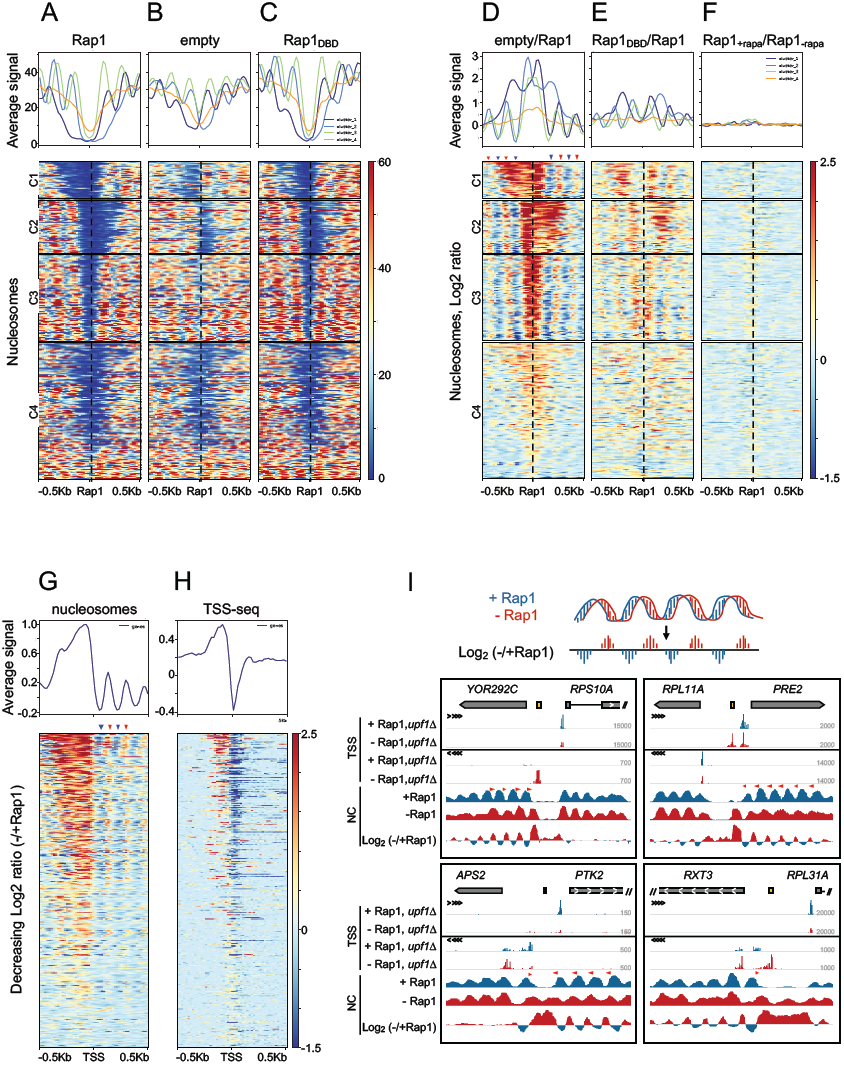
Nucleosome and TSS selection changes upon depletion of Rap1 and expression of the Rap1 DNA binding domain alone (Rap1_DBD_). See also Figure S3. A-C : heatmaps showing the profile of MNase-seq signals at genes containing a Rap1 site upon nuclear depletion of Rap1 and ectopic expression of a wild type gene (A), an empty vector (B) or a construct containing only the Rap1 DNA binding domain (C). D-F : heatmaps reporting the Log_2_ratio of the MNase-seq signal change upon depletion of Rap1 and expression of the empty vector (D) or the Rap1_DBD_(E). F is a control for the signal change upon expression of ectopic Rap1 under the same conditions as D and E. Genomic regions aligned on Rap1 sites and sorted by k-means clustering. Metanalysis for each cluster are shown on top. G: heatmap showing the MNase-seq signal change (Log2 ratio) upon Rap1 depletion at genes aligned on their TSS. Features are sorted by decreasing signal change. Alternate stripes of red and blue downstream of the alignment point (red and blue arrowheads) indicate changes in the phasing of the genic nucleosome array at many genes when Rap1 is depleted (see also I) H: Genes are aligned and sorted as in G, but the Log_2_ratio of the TSS-Seq signal change upon Rap1 depletion is shown, indicating that changes in TSS usage strongly correlate with nucleosome positioning changes. I. Individual examples of correlations between changes in the pattern of initiation and alterations of the MNase-seq profile. The movement of nucleosomes in Rap1-depleted cells (red tracks) is indicated by small red arrowheads in the wild type tracks (blue) and by the characteristic sigmoidal pattern of the Log_2_ratio due to nucleosome dephasing (see scheme on top). The latter is responsible for the “striped” pattern seen in G.

In many cases we also observed the appearance of non-coding transcription running antisense to a Rap1 gene target, indicating that Rap1 restricts the intrinsic bidirectionality of many promoters and limits pervasive transcription at the level of initiation (Figure 1 J-K and Figure S1K-L). To assess the generality of this finding we aligned all the Rap1 sites that lack an annotated coding gene within the downstream 500nt and profiled the log2 ratio of the RNAPII CRAC signals observed in the absence or presence of Rap1. As shown in fig. 1L, many novel, mostly non-coding, transcription events are generated when Rap1 is absent from the nucleus, often initiating immediately downstream of the Rap1 site. Visual inspection of the regions listed in figure 1L (304 sites) allowed estimating that Rap1-suppressed non-coding transcription occurs in roughly 30% of cases (see below).

Finally, alterations in gene expression were not exclusively associated to ectopic initiation, some genes demonstrating decreased transcription initiation from canonical sites in the absence of eTSS (Figure S1M-O).

Together, these results demonstrate that Rap1 represses non-coding and ectopic transcription initiation in promoters. Many novel transcripts are synthesized extremely rapidly after Rap1 depletion, underlying the appearance of an unexpectedly complex landscape of gene expression effects.

### Diverse effects of Rap1 depletion on gene expression

Ectopic initiation can have multiple consequences on gene expression. Transcription initiated at ectopic sites (eTSS) can invade the downstream gene and inhibit or overlap transcription initiated at the canonical site. The novel transcripts generated might code for different protein isoforms or contain premature stop codons targeting them to nonsense-mediated decay in the cytoplasm. Finally, some of these novel transcripts can be subject to quality control by the exosome and degraded rapidly after transcription in the nucleus.

To disentangle overlapping transcription events derived from different initiation sites we mapped by TSS-Seq (Malabat et al., 2015) the 5’ends of transcripts produced after nuclear depletion of Rap1 for 60 minutes, which we estimated to be a good compromise for reliably detecting eTSS appearance while minimizing secondary effects. To detect transcripts that might be unstable, we also analyzed TSSs in cells defective for either nuclear degradation (*rrp6Δ*) or nonsense-mediated decay (*upf1Δ*). The data obtained revealed a wealth of novel RNA species arising from ectopic transcription. Upon depletion of Rap1, spurious initiation most often occurs in the vicinity of canonical initiation sites (Figure 2 A-C, G-I) and in some instances at or close to the site of Rap1 binding (Figures 2G, I, K; Figure S2 A-B; see also Figure 1N). Many of these initiation events generate RNA species that are unstable and at least partially sensitive to NMD (Figure 2D; 2G-I, compare –Rap1, *upf1Δ* with –Rap1 tracks) or nuclear degradation (Figure 2E, 2K). No spurious initiation was observed upon rapamycin treatment of cells not expressing Rap1-FRB (Figure 2F) or at genes not containing Rap1 binding sites in the promoter region (compare Figure S2C and S2D). A marked downregulation at canonical initiation sites was observed at many Rap1 targets (Figure 2A-C), often in association with ectopic initiation, mirroring the downregulation observed by RNAPII CRAC (Figure 1A-F).

The case models described below illustrate the variegated and sometimes complex impact on gene expression ensuing from this deregulation of TSS selection.

### Ectopic initiation and downregulation of gene expression

The paradigmatic case of the *PRE2* gene is shown in Figure 2G. The RNAPII CRAC signal arising upstream of the gene upon Rap1 depletion generates a transcript starting roughly 140nt upstream of the canonical *PRE2* TSSs. This RNA molecule contains two upstream ORFs (top scheme, red arrow), which explain its marked sensitivity to nonsense-mediated decay (Figure 2G, compare tracks –Rap1 and –Rap1/*upf1Δ;* Figure 2L, lanes 3 and 6). Northern blot analysis (Figure 2L) confirmed that this species is longer than the canonical *PRE2* RNA suggesting that it shares the same 3’-end. The appearance of this novel 5’-extended RNA is accompanied by a marked decrease in transcription starting at the canonical *PRE2* TSS (Figures 2G, inset, and 2L, compare lanes 1 and 3). Consistently, the levels of a FLAG tagged form of the Pre2 protein (Figure 2M) were markedly reduced in coincidence with the TSS shift. Thus, lower expression from the canonical TSS and production of an unstable RNA explain downregulation of *PRE2* expression in the absence of Rap1. Note that the transcription signal observed in the body of *PRE2* is not a faithful predictor of *PRE2* expression as it corresponds to production of mRNA species only some of which have the right coding potential. Cases akin to *PRE2* are relatively frequent among Rap1 gene targets (15-20% of the total), notably genes coding for ribosomal proteins (Figure S2E-F).

### Apparent upregulation of gene expression

Besides cases of downregulation, we observed apparent gene up-regulation upon selection of an upstream eTSS. One example is shown in Figure 2H for the *NAT3* locus (see also *YOR292C*, Figure 3I). In the absence of Rap1 an upstream TSS is selected to levels markedly higher than the natural TSS, leading to increased transcription levels from an altered start site as demonstrated by the RNAPII CRAC signal. A consistent fraction of these RNAs are degraded in the cytoplasm by the NMD pathway as they contain upstream ORFs (uORFs) (Figure 2H), but the molecules that escape degradation are still more abundant than the correctly initiated *NAT3* RNAs. Thus, although both transcription and steady state RNA levels for *NAT3* (and *YOR292C*) appear to be upregulated in the absence of Rap1 (potentially qualifying the latter as a repressor) the RNAs produced do not have the same coding potential as the RNA initiated at the canonical TSS.

An interesting variant of apparent upregulation is represented by the *RTX3* locus (Figures 2I and 2L). In this case, transcription is naturally started from two TSS clusters, one that is internal to the ORF, and a second upstream of the natural ATG. Use of the internal site leads to the production of a truncated protein, lacking the first 51aa. Depletion of Rap1 induces strong repression of the natural TSS that is accompanied by strong induction of the internal TSS and by the selection of an additional upstream eTSS generating NMD-sensitive transcripts. This leads to an overall increased transcriptional and steady state RNA signal associated with the *RXT3* locus. Still, this corresponds to increased production of a truncated isoform (Figure 2M, red arrow) and to decreased levels of the normal protein.

These results demonstrate that many cases of apparent transcriptional upregulation of gene expression in the absence of Rap1 in reality hide a constellation of scenarios that generally converge on downregulation of functional transcripts.

### Bona fide upregulation following Rap1 depletion

Alterations in TSS usage in the absence of Rap1 can also lead to a *bona fide* increase in gene expression. Although not frequent, the case of *APS2* nicely illustrates this situation (Figures 2J, 2L-M). In the presence of Rap1 transcription initiates at two sites: the ATG proximal site is responsible for the expression of the functional *APS2* RNA, whereas a preferentially used distal site leads to the expression of a transcript that contains premature stop codons and is subject to NMD (Figure 2J; 2L, compare lanes 1 and 4). Upon Rap1 depletion, the ATG-proximal site is favored over the upstream site (Figures 2J, 2L, lanes 3 and 6), leading to the increased production of a stable *APS2* RNA and a functional Aps2 protein (Figure 2M). In this case, Rap1 functions as a *bona fide* repressor of *APS2* expression. However, repression occurs by a non-canonical mechanism involving a reorganization of TSS usage that favors a functional site.

### Inhibition of the canonical TSS usage is not due to transcriptional interference

It has been shown that when transcription occurs within a promoter in the sense or antisense direction it can inhibit its function by establishing a repressive chromatin structure (for reviews see Jensen et al., 2013; Porrua and Libri, 2015). Therefore upstream transcription from the eTSS might inhibit downstream initiation because of transcriptional interference. Alternatively, decreased use of the canonical TSS (e.g. at the *PRE2* locus) might occur independently from the occurrence of upstream transcription, for instance as a consequence of altered TSS selection preferences in Rap1-deficient cells.

The histone methyltransferase Set2 and the histone deacetylase SET3 complex have been shown to be essential for transcription interference (Kim et al., 2016, 2012). We therefore depleted Rap1 in a *set2Δ* or a *set3Δ* context, and in the absence of Upf1 to improve detection of unstable RNA species. Northern analysis of the RNAs derived from the *PRE2, RXT3* and *APS2* loci did not reveal restoration of the downstream TSS usage in the absence of Set2 or Set3 (Figure 2L; note that in the case of *APS2* inhibition of the downstream TSS is expected to occur in the presence of Rap1). We conclude that, at least at the loci analysed, repression of downstream TSS usage does not results from Set2 or Set3-dependent transcription interference.

### Altered nucleosomal architecture underlies spurious transcription initiation

We next investigated the mechanism by which Rap1 controls the fidelity of transcription initiation. It has been shown that depletion of Rap1 is linked to the appearance of Micrococcal Nuclease (MNase)-resistant segments in many NDRs, proposed to represent “fragile nucleosomes” that in the presence of Rap1 are only detected when using low MNase concentrations (Knight et al., 2014; Kubik et al., 2015). Increased nucleosome occupancy (or decreased MNase accessibility) in NDRs is generally linked to inhibition of transcription initiation, which is seemingly at odds with the appearance of novel TSS. Therefore, we analysed the impact of Rap1 on the nucleosomal architecture in and around the NDR to the light of our results. Because we also analyzed the position of nucleosomes in cells expressing a variant of Rap1 (see below) we produced novel MNase-Seq data upon Rap1 nuclear depletion. As shown in Figure 3A,B,D (regions aligned on Rap1 sites and sorted by K-means clustering) and Figure S3A,B,D (regions aligned on TSSs downstream of Rap1 sites and sorted by K-means clustering), depletion of Rap1 markedly increased overall nucleosome occupancy in roughly half of the NDRs bound by Rap1, consistent with previous reports (Knight et al., 2014; Kubik et al., 2015). A closer look at the differential MNase resistance signal also revealed a striped pattern (blue and red arrowheads in Figures 3D and 3G) indicating that in many instances at least part of the whole nucleosomal array bordering the NDR is phase shifted relative to the wild type pattern and moves towards the unoccupied Rap1 site (see individual cases in Figure 3I; note the characteristic sinusoidal behavior of the log2 ratio of nucleosome occupancy). Thus, increased nucleosome occupancy in Rap1-dependent NDRs depends on the frequently observed +1 nucleosome shifting, on the addition of an extra nucleosome or both.

When we ordered Rap1 sites based on nucleosomal occupancy change (Figure 3G), we observed clear correlation with changes in TSS usage (both emergence of eTSS and decreased use of normal TSS, Figure 3H), which strongly suggests that the altered pattern of nucleosome positioning in and around the NDR is the cause of the massive effects on transcription initiation induced by the absence of Rap1. This is clearly illustrated by the paradigmatic examples shown in Figure 3I (see also Figure S3G).

The upstream shift of four to six nucleosomes at the *YOR292C* and *PRE2* loci is associated to an upstream initiation shift that occurs at the site that was previously occupied and presumably sterically hindered by Rap1. At the *APS2-PTK2* divergent locus, the concomitant upshift of one nucleosome on the *APS2* side and 5-7 nucleosomes on the *PTK2* side dramatically shrinks the main wild type NDR and opens an alternative NDR between the original *APS2* +1 and +2 nucleosomes. This hinders the main, non-functional *APS2* TSS (now buried under the shifted +1 nucleosome) and favors use of the site that generates a functional transcript (Figure 2M). An additional eTSS can also be detected upstream of *PTK2*

The complex transcription initiation pattern observed at the *RXT3* locus can also be explained by local changes in the nucleosomal architecture (Figure 3I). The upstream shift of the +1 nucleosome can be accounted for hindering the canonical TSS, inducing an upstream eTSS and creating a small NDR between the +1 and +2 nucleosomes that favors the use of the internal TSS coding for a N-ter truncated isoform (Figure 2M).

A similar case in which expression of N-ter truncated isoforms is induced is shown for the *PEX12-YMR027W* locus in Figure S3G. At this interesting locus, the upstream shift of 4 *PEX12* nucleosomes is associated with internal initiation between the fourth and fifth nucleosomes, indicating that the absence of Rap1 can induce ectopic initiation at a distance. Upshift of the +1 nucleosome also induces internal initiation at the divergent *YMR027W* gene, with strong downregulation of the normal TSS.

Together, the genome-wide analyses and the single examples illustrate a strong correlation between the modified nucleosomal architecture observed in the absence of Rap1 and the massive alterations in the selection of the transcription initiation site. Upstream shifts of +1 nucleosomes generally lead to TSS downregulation. However, downregulation of canonical initiation is often associated with the emergence of new transcription initiation events that generally arise in close correspondence to the newly positioned nucleosomes.

### The DNA binding domain of Rap1 can support normal nucleosome positioning and transcription initiation

The mechanism of Rap1 action on nucleosomes and on gene activation/repression is still not well understood in spite of a wealth of studies on the matter. Rap1 interacts directly with the SWI/SNF chromatin remodelling complex (Tomar et al., 2008) and has been shown to be required for the recruitment of Esa1, a histone H4 acetylase (Reid et al., 2000). Activation of the expression of model genes by Rap1 depends on a domain that is C-terminal to the DNA binding domain (DBD) (Garbett et al., 2007; Johnson and Weil, 2017; Papai et al., 2010; Tomar et al., 2008), which has been implicated in the interaction with TFIID and SWI/SNF (Johnson and Weil, 2017). To assess whether the role of Rap1 in controlling the fidelity of initiation is related to the recruitment or function of TFIID or SWI/SNF, we depleted Rap1 from the nucleus of cells ectopically expressing only the DBD of Rap1 (aa. 358-601, Rap1_DBD_), an empty vector or the wild type protein were used as controls. Under these conditions, expression of Rap1_DBD_ did not support viability and affected normal yeast growth even in the presence of wild-type Rap1 (data not shown). Rap1_DBD_ binds DNA *in vivo*, because it could induce transcription termination at a site where Rap1 roadblocks RNAPII (Candelli et al., 2018).

The growth defects induced by expression of Rap1_DBD_ precluded reliable RNAPII CRAC analyses of these strains as well as the mutation of degradation pathways. However, in spite of their sensitivity to NMD, many 5’-extended transcripts produced in the absence of Rap1 could still be detected in otherwise wild type cells (Figure 2 and 4). Therefore we restricted our analyses to these RNA species as a proxy for the occurrence of ectopic initiation and analysed in parallel the MNase sensitivity profile in cells expressing Rap1_DBD_ after transient depletion of wild type Rap1. To our surprise, Rap1_DBD_ suppressed to a very significant extent the nucleosomal phenotype of Rap1-deficient cells (Figures 3A-D and Figures S3A-D). This indicates that the maintenance of a quasi-normal nucleosomal architecture at many Rap1 targets does not require domains of Rap1 involved in transcriptional activation (Johnson and Weil, 2017). Interestingly, however, suppression was not complete (compare Figures 3A to 3C and Figure 3E) and was most prominently observed in the region immediately surrounding the Rap1 binding site (Figure 3E and Figure S3E). Such a different chromatin architecture at the same sites upon expression of Rap1DBD provided an unexpected opportunity for further correlating ectopic initiation with the altered position of nucleosomes.

**Figure 4:**
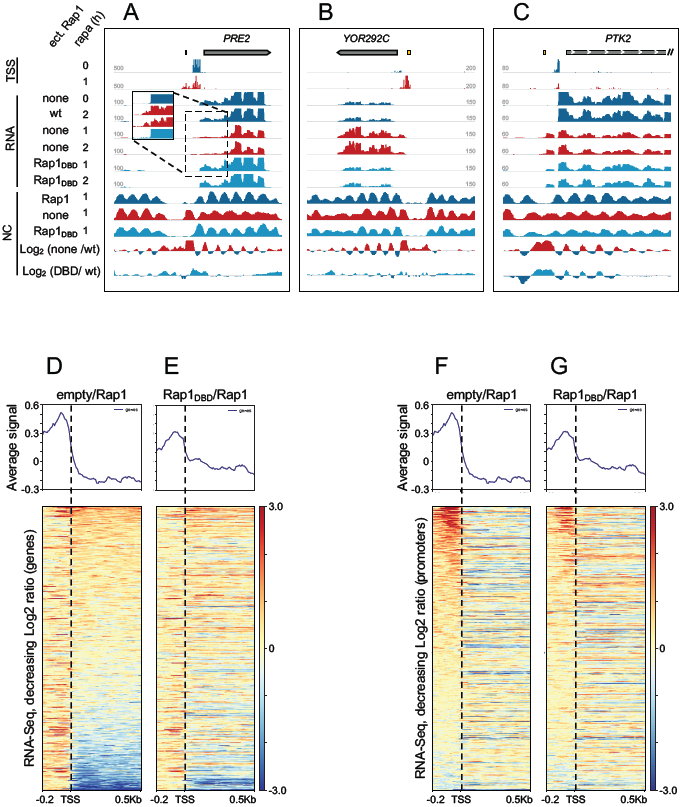
Rap1_DBD_ partially suppresses the gene expression phenotypes linked to Rap1 depletion. See also Figure S4. A-C: Individual snapshots are shown to illustrate cases of full (A,B) or poor (C) suppression, which correlate with nucleosome positioning changes. Additional examples are shown in Figure S4. D-G. Heatmaps showing the RNAseq signal change (Log2 ratio) upon Rap1 depletion in the absence of ectopic Rap1 expression (D, F) or in the presence of Rap1_DBD_ (E, G). Genes are sorted according to decreasing signal change in in the first 500nt of genes D,E) or in the promoter region (−200 to +5 relative to the TSS, F,G) to illustrate the extent to which Rap1_DBD_ suppresses downregulation of Rap1 target genes (E versus D) and ectopic TSSs selection (G versus F). Note that only the fraction of ectopic initiation events producing transcripts that survive NMD or nuclear degradation are detected in this analysis. A statistical analysis of the suppression is shown in Figure S4C-E.

Individual snapshots correlating nucleosome positioning and transcripts produced in the presence of Rap1_DBD_ are shown in Figure 4A-C and Figure S4A-B.

At the *PRE2* and *YOR292C* loci, Rap1_DBD_ restores an apparently normal position of nucleosomes as well as the normal site of transcription initiation and the level of average steady state RNA (which is higher in the wild type for *PRE2* and lower for *YOR292C*). Conversely, nucleosome positioning is not restored at the *PTK2* locus, and ectopic initiation leading to the expression of a 5’-extended RNAwas maintained upon expression of Rap1_DBD_. Within large NDRs with an eccentrically positioned Rap1 site, expression of Rap1_DBD_ only restores normal position of nucleosomes and transcription initiation around the binding site (for two examples see Figure S4A-B).

For a genome-wide assessment of the extent of the Rap1_DBD_ suppression, we sorted all Rap1 targets genes by RNAseq signal change within genes (Log_2_ratio) upon Rap1 depletion (Figure 4D). This provides an overall view of gene deregulation based on RNA abundance. Analysis of the same signal in the presence of Rap1_DBD_ clearly indicates that Rap1_DBD_ restores the normal levels of many (although not all) transcripts (Figure 4E). When we considered all the transcripts whose levels were decreased by at least a factor of 2 in Rap1-deficient cells (roughly 30% of the total), restoration by Rap1_DBD_ could be observed at a high level of significance (p=2*10^−6^; Supplementary Figure 4C). To focus on ectopic initiation events, we sorted the same set of genes by decreasing signal change upon Rap1 depletion in the 200 nt upstream of the TSS (Figure 4F). The increased RNAseq signal present in the promoter of these RNAs was partially restored in the presence of Rap1_DBD_ (Figure 4G and Figure S4D; p=2,7*10^−5^). We also assessed the statistical significance of the suppression of non-coding transcription by Rap1_DBD_ We computed the increase in RNAseq signal around Rap1 sites that are not followed by mRNA-coding genes within 500 nucleotides (same set as in Figure 1L). At the sites showing the highest levels of non-coding transcription increase (57), suppression by Rap1_DBD_ was again statistically significant (p=6.5*10^−4^, Figure S4E). Note that many non-coding RNA ectopic initiation events are not detected in NMD and exosome proficient cells. Finally, no effect was observed at unrelated genes (Figure S4F).

Together these results demonstrate that expression of only the DNA binding domain of Rap1 is sufficient to restore the canonical position of nucleosomes at many sites in and around the NDR. Suppression of the nucleosomal architecture is not always complete, but the correlation between nucleosome positioning and the altered usage of ectopic and canonical TSSs is strikingly maintained.

### The control on initiation fidelity by Rap1 is not mediated by SWI/SNF, RSC or INO80 chromatin remodeler complexes

Although Rap1_DBD_ can partially suppress nucleosome positioning phenotypes due to the absence of Rap1, we cannot formally exclude that Rap1 displaces nucleosomes by recruiting chromatin remodelers. Possibly consistent with this notion, it has been shown that SWI/SNF still interacts with Rap1_DBD_, albeit to a lesser extent than with the wild type protein (Tomar et al., 2008). It has also been proposed that the RSC chromatin remodelling complex functions downstream of the Reb1 and Abf1 GRFs at many NDR (Hartley and Madhani, 2009). If chromatin remodelers were downstream effectors of Rap1 action, affecting their function should lead to a similar and possibly more general phenotype as depletion of Rap1. We therefore assessed the effect on transcription initiation upon depletion of the catalytic subunits of the SWI/SNF or RSC complexes, Snf2 and Sth1 respectively. We also co-depleted Isw2 and Ino80, which are ATPases involved in chromatin remodelling that have been implicated, possibly redundantly, in the positioning of the +1 nucleosome (Krietenstein et al., 2016; Whitehouse et al., 2007). The comparative effects on nucleosome depletion of these factors will be discussed in detail in a separate report (Kubik et al., submitted), we will present here only results relative to Rap1 targets.

Transcription start sites were mapped genome-wide after single depletion of Sth1 and Snf2 or concomitant depletion of Ino80 and Isw2. As previously for Rap1, we also performed the same experiments in an *upf1*Δ background. The distribution of TSSs at Rap1 targets sorted according to decreasing levels of MNase seq signal changes in Rap1-deficient cells is shown in Figures 5B-E. Individual snapshots are shown in Figures 5F-I, where we also show the changes in nucleosome occupancy in these conditions (Kubik et al., submitted). The levels and distribution of ectopic initiation were strikingly different in Sth1, Snf2 or Ino80/Isw2 deficient cells compared to cells lacking Rap1. Although novel TSSs were observed in NDRs upon Sth1 and Snf2 depletion, their distribution was qualitatively different than that of Rap1-repressed eTSSs, which strongly argues against a common mechanism (Figure 5F-I). Of note, upon co-depletion of Ino80/Isw2 the pattern of ectopic transcription initiation was complementary and less defined relative to that observed in Rap1-depleted cells, with eTSSs frequently present within genes and in the same direction of transcription, presumably due to the less well-positioned intragenic nucleosomes (Figure 5).

**Figure 5:**
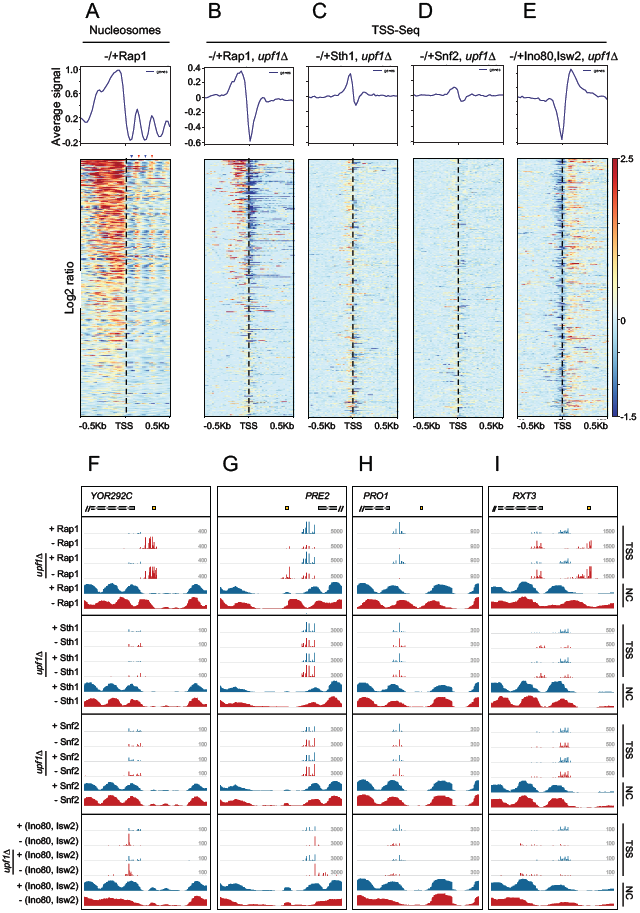
TSS usage changes upon depletion of chromatin remodelers compared to changes upon depletion of Rap1. A-E : in all heatmaps features are sorted according to changes in MNase-Seq signal upon Rap1 depletion (A) as in Figure 3G. Changes in TSS usage (Log_2_ratio) upon depletion of the indicated chromatin remodeler subunits are shown in C-E and compared to signal changes upon Rap1 depletion (B, same as Figure 3H, shown here for ease of comparison). Only signals derived from cells defective for NMD (*upf1Δ*) are shown, but similar results have been obtained with NMD proficient cells. F-I snapshots comparing TSS and nucleosome changes upon depletion of Rap1 or the indicated chromatin remodeler subunits. Data from depleted cells are shown in red. The position of the Rap1 site is indicated.

From these experiments we conclude that Rap1 suppresses ectopic transcription initiation independently of chromatin remodelers that might be recruited at its target sites. These results also suggest that the two pathways act synergistically for promoting transcription initiation fidelity, most likely because of their cooperation in nucleosomes positioning (Kubik et al., 2018)

### Reb1 and Abf1 suppress ectopic transcription initiation

Reb1 and Abf1 are GRFs that have similar roles to Rap1 in terms of nucleosome exclusion at NDR (Badis et al., 2008; Ganapathi et al., 2011; Hartley and Madhani, 2009; Kubik et al., 2015). We analysed the role of both factors in suppressing ectopic initiation at their target genes by profiling RNAPII occupancy by CRAC upon nuclear depletion of Reb1 or Abf1. Depletion of either factor led to increased nucleosome occupancy in NDRs (Figures 6A and 6E, data from Kubik et al., 2015), as previously reported, and shown for Rap1 in Figure 3. We also observed a very frequent upstream shift of the intragenic nucleosomal array relative to the canonical TSS, as observed for Rap1 depletion, responsible for the characteristic “striped” pattern in the differential signal (Figures 6A and 6E).

**Figure 6:**
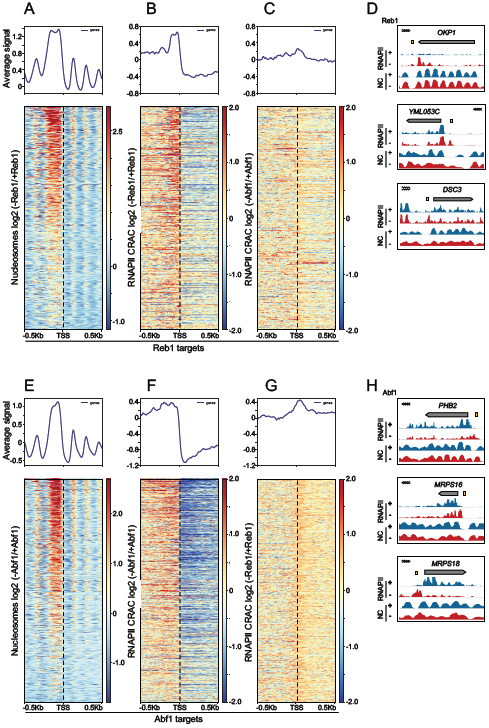
Nucleosome occupancy and TSS selection changes upon depletion of Reb1 and Abf1. A. TSSs of genes containing a Reb1 site within the upstream 300nt (650) were aligned and sorted according to decreasing nucleosome changes. B. RNAPII CRAC signal changes upon Reb1 depletion. The same set of genes are aligned to canonical TSSs and sorted as in A.C. same as B, but the RNAPII CRAC signal change in Abf1-deficient cells is plotted instead of Reb1-deficient cells (note that this is a specificity control for Abf1 depletion. The control for Reb1 depletion is shown in G). D. Individual snapshots illustrating the occurrence of ectopic initiation at Reb1 targets upon Reb1 depletion. In the top panel, depletion of Reb1 induces non-coding transcription antisense to *OKP1.* In the bottom panels, the absence of Reb1 is associated to changes in nucleosome positioning and ectopic initiation upstream of *YML053C* and *DSC3*.E. TSSs of Abf1 gene targets (781) are aligned and sorted according to decreasing nucleosome changes as in A. F. RNAPII CRAC signal changes at Abf1 target genes upon Abf1 depletion as in B. G. same as F, but the changes in RNAPII CRAC signal upon Reb1 depletion is shown as a control for the effect of Reb1 depletion at unrelated genes. Note that in all heatmaps genes that contain both a Reb1 and Abf1 binding site have been excluded.

**Figure 7:**
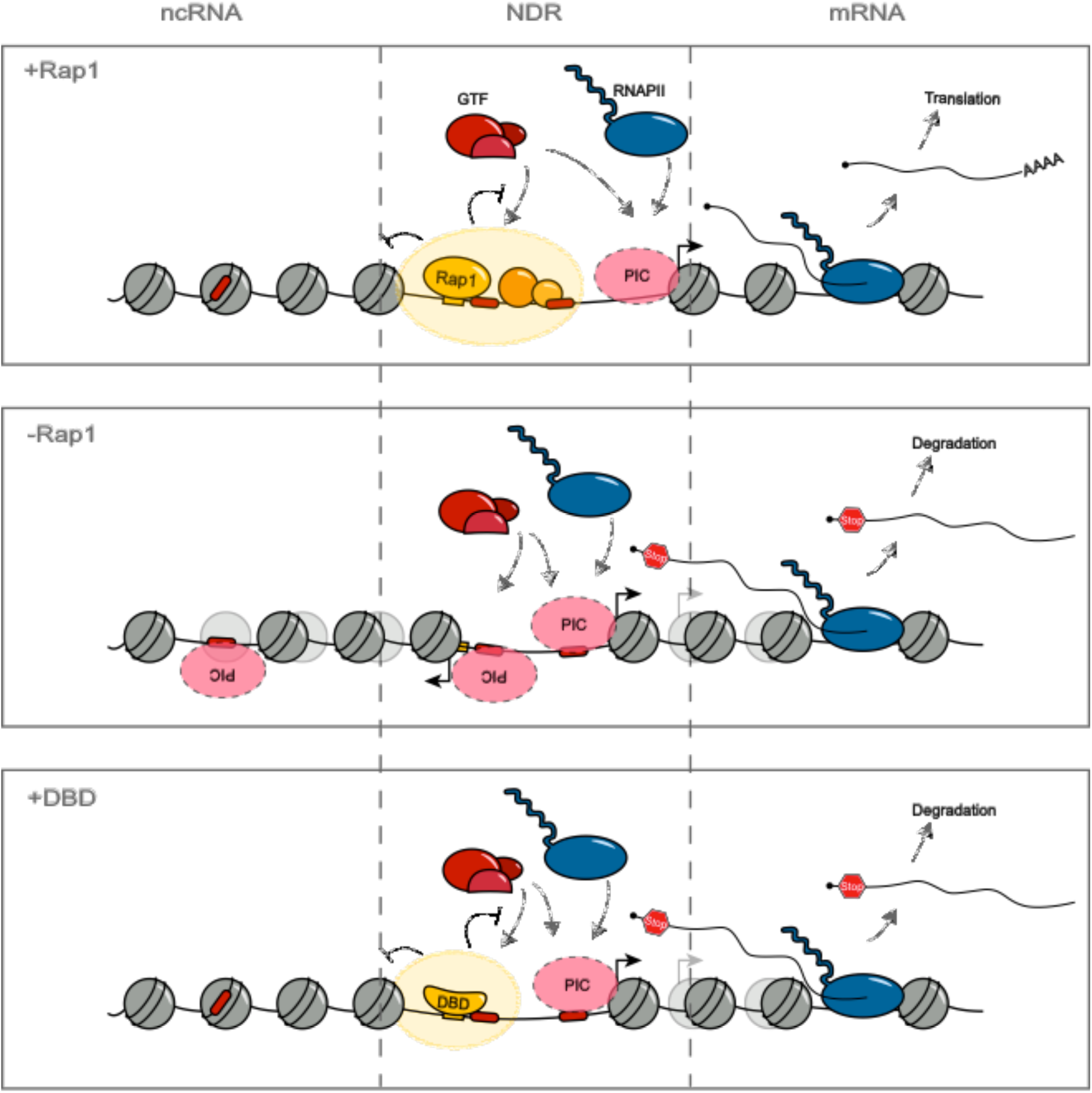
Model illustrating the role of Rap1 and GRFs in the suppression of ectopic initiation. A. In wild type cells Rap1 (and other GRFs) participate to the positioning of nucleosomes that are excluded from the NDR, at least in part by a steric mechanism. In the case of Rap1 the associated FIS (Fhl1, Ifh1 and Sfp1) and Hmo1 could also be required for excluding nucleosomes from the NDR. General transcription factors (GTFs) could access the DNA at cryptic sites (red boxes) and generate mispositioned initiation events, which are however prevented by the bound Rap1 and other GRFs (yellow ellipse). In the absence of Rap1(B) these cryptic PIC formation sites become accessible, leading to the production of non-coding (leftward) or miscoding RNAs (rightward). The novel positioning of nucleosomes that encroach the NDR leads to extensive alterations in the definition of the +1 nucleosome for initiation. C. When the sole DNA binding domain of Rap1 is expressed (DBD), in many instances nucleosome positioning is restored, mainly in the vicinity of the site, together with the suppression of cryptic PIC formation and eTSSs. In this example we show one of the large, Rap1-dependent, NDRs in which restoration only occurs on the side of the Rap1 binding site (yellow box).

Heatmaps reporting the changes in the RNAPII CRAC signal upon depletion of Reb1 and Abf1 (Figures 6B and 6F respectively) clearly demonstrate the occurrence of transcription initiation in the NDR upstream of Reb1 and Abf1 targets, which is generally associated with downregulation of the downstream gene (see also the summary plots associated to each heatmap). As a control, Reb1 and Abf1 depletion did not induce ectopic initiation at the non-cognate targets (respectively Abf1-dependent, Figure 6G, and Reb1-dependent, Figure 6C). As for Rap1, changes in nucleosome positioning induced by the absence of Reb1 or Abf1 strongly correlate with the appearance of novel eTSSs and the downregulation of the canonical site of initiation. In many instances, we observed effects on gene expression that were not previously noticed based on steady state RNA level changes, most likely because they were masked by the overlapping with 5’-extended transcripts (data not shown).

Together these results extend the essential role of Rap1 in controlling the fidelity of transcription initiation to other GRFs and possibly to other transcription factors.

## DISCUSSION

In this study we have addressed the effects on transcription and on gene expression of depleting Rap1 and two other GRFs, Abf1 and Reb1. These factors have been known for decades to affect the expression of many highly expressed genes. Still, their mode of action for gene activation or repression, has remained relatively obscure. Here we analyzed transcription directly by RNAPII CRAC and complemented these data with the assessment of transcription initiation sites of stable and unstable transcripts upon rapid depletion of these factors. Overall, we discovered that transcription of target genes was not generally shut off in the absence of these proteins, as one would expect for typical trans-activators. Rather, the landscape of transcription initiation was radically modified, revealing a myriad of novel transcripts and isoforms that alter in a manner that is not predictable a priori the ultimate outcome of gene expression. These effects are related to the modified positioning of nucleosomes in the NDR and also in the body of genes, but are distinct from what observed when individual or combination of chromatin remodelers are depleted. Importantly, the sole DNA binding domain of Rap1 is sufficient for full restoration of nucleosome positioning and repression of ectopic initiation at many sites, pointing to a “place-holder” model for the impact of Rap1 (and possibly other GRFs) on the chromatin architecture of NDRs.

Our findings imply that Rap1 and other GRFs control the fidelity of transcription initiation, preventing deleterious and non-coding transcription events from taking place within the nucleosome depleted regions to which they bind. A parallel study from the van Werven laboratory reached similar conclusions (Wu et al., 2018).

### Rap1 and nucleosome positioning

Many earlier studies have shown that Rap1 is important for the size and nucleosome occupancy of the NDRs to which it binds (Badis et al., 2008; Ganapathi et al., 2011; Hartley and Madhani, 2009; Kubik et al., 2015), which we also have observed. One important facet of our work is the demonstration that the sole DNA binding domain of Rap1 is sufficient for restoration of normal nucleosome positioning and TSS selection fidelity at many sites of Rap1 binding. We also show that the strong effect on nucleosome positioning (and TSS selection) due to the absence of Rap1 cannot be ascribed to defective recruitment or function of chromatin remodelers, at least the ones we tested, i.e. the RSC, the SWI/SNF, Isw2 and Ino80 complexes. This suggests that Rap1 constrains nucleosomes at least in part by a steric hindrance mechanism, which is also consistent with suppression by Rap1_DBD_ occurring most efficiently around the Rap1 binding site (Figure 3E and Figure S3E). Earlier results from the Morse lab showing that Rap1 variants lacking either the N- or the C-terminal domains of the protein can alter the nucleosomal structure of a reporter construct (Yu et al., 2001) are fully consistent with our findings. *In vitro* reconstitution experiments with purified factors have shown that GRFs are not intrinsically required for NDR formation, but are important for positioning the +1 nucleosome in the presence of ISW2/ISW1 remodelers (Krietenstein et al., 2016). Although Rap1 was not used in these experiments, it is possible that like other GRFs it constitutes a physical barrier against which genic nucleosomes are “pushed” by ISW2/ISW1, a “place-holder” function for which in many instances only DNA binding is absolutely required. We observed, however, that when the Rap1 binding site is eccentric relative to the NDR, Rap1_DBD_ can only constrain proximal nucleosomes, which implies that the domains missing in Rap1_DBD_ are important for maintaining the integrity of the NDR at some distance from the binding site. The reason for this is unclear, but might be related to the interaction of factors frequently associated to Rap1 in the NDR, such as Fhl1/Ifh1, Sfp1 (FIS) and Hmo1 (Knight et al., 2014; Reja et al., 2015). This notion is fully consistent with earlier results showing that the absence of Hmo1 induces an upstream shift of the +1 nucleosome that inhibits gene expression (Kasahara et al., 2011; Reja et al., 2015) and would imply that components of the FIS and/or Hmo1 are not bound, or cannot restrict nucleosome displacements when Rap1_DBD_ is bound to the NDR.

### The chicken and egg issue of nucleosomes and transcription initiation

In yeast transcription initiation is known to occur at 12-15 nt of the upstream border of the +1 nucleosome (Hughes et al., 2012; Lee et al., 2007; Tsankov et al., 2010; Whitehouse et al., 2007). Whether it is initiation that specifies the exact position of the +1 nucleosome or the latter that directs the position of initiation is still to some extent a matter of debate (for recent reviews see Lieleg et al., 2015; Struhl and Segal, 2013). Once the PIC is established, RNAPII scans the downstream sequence for the most favorable site of initiation, which is generally located at least 50nt downstream. Although the position of the TSS is also defined by its surrounding sequence, the consensus established is loose (Malabat et al., 2015) and unlikely to be the sole determinant. We observed a very close relationship between the positions of nucleosomes and ectopic initiation, which invariably coincide in the sense that eTSSs are always found on the upstream border of a displaced nucleosome. Even subtle upstream displacements are sufficient to shift the pattern of initiation, an observation that also holds for chromatin remodelers (see for instance the effect of depleting Sth1 and Snf2 at the *YOR292C* locus, Figure 4). Interestingly, this rule is also met when the nucleosomes shift downstream as in the double depletion of Ino80 and Isw2: in these instances (Figure 4), the canonical TSS remains exposed to the scanning RNAPII, but it is not (or poorly) used, presumably because it is incorrectly positioned relative to a nucleosome. Although it could be argued that these factors affect primarily the position of transcription initiation and the position of the +1 nucleosome as a consequence, the most parsimonious explanation for these findings is that the position of the +1 nucleosome causally underlies TSS selection. Whether initiation determines the peculiar composition of the +1 nucleosome (e.g. the presence of H2AZ) or whether the latter preexists and is one determinant of initiation remains matter of speculation.

### Rap1 controls the fidelity of transcription initiation

The absence of Rap1 brings about a major upheaval in the nucleosomal landscape in and around the NDR. We expected that this would cause extensive downregulation of target genes, linked to the frequent upshift of the +1 nucleosome (Parnell et al., 2015; Reja et al., 2015; Shivaswamy and Iyer, 2008; Whitehouse et al., 2007). What was less expected, however, is that increased nucleosome occupancy in the NDR would allow such a massive number of novel initiation events, many of which are non-coding or generate RNAs with lower/altered coding potential. It thus appears that whenever the new eTSS is not positioned within the PIC-TSS RNAPII scanning window (on average 50-100nt), this leads to the ectopic formation of novel PICs, which drive transcription in the sense or antisense direction relative to Rap1- (and other GRF-) controlled genes. The appearance of eTSSs was indeed invariably associated with the presence (or formation) of an upstream MNase-sensitive region. Because there is no reason to hypothesize the existence of some evolutionary pressure to maintain these ectopic PICs, it must be concluded that they form promiscuously in the absence of a specific negative control mechanisms. Since stable nucleosomes are absent from these regions in wild type cells, they cannot be accounted for hindering formation of ectopic PICs at these sites. Therefore, the role that Rap1 and other GRFs fulfill in preventing formation of these PICs under normal conditions is likely independent from their role on nucleosome positioning and we suggest that is linked to the GRF and GRF-dependent physical occupancy of these sites. However, we cannot exclude that the presence of nucleosomes is actually *required* for transcription initiation to take place. For instance it could be imagined that once a PIC is formed, a nearby nucleosome is required for transcription initiation, otherwise the PIC would not be productive. Or, alternatively, that a nearby nucleosome could favor formation of a PIC.

The quality control on the fidelity of initiation fulfilled by GRFs is expected to be structural and constitutively required as its absence has major and disruptive effects as shown here. Its modulation, could be, however, a powerful tool for gene regulation (Bosio et al., 2017; Reja et al., 2015).

### Impact of novel TSSs on gene expression

Another important implication of this study is that neither the assessment of RNAPII levels nor the steady-state levels of RNA within genes can be used as a reliable proxy for inferring gene expression levels, unless information concerning the position of the TSS is also integrated into the equation. This implies that the notion of positively or negatively regulated genes has to be generally revisited, at least for the GRFs that we have studied, but potentially for all of the factors that might affect the selection of transcription initiation sites or the position of nucleosomes. We found several examples in which overlapping transcription signals initiated at a canonical and an ectopic TSS leads to an apparent increase in gene expression, or an apparent lack of effect. However, in the majority of cases deconvolution of the two signals using TSS analysis demonstrates a likely decrease in the ultimate outcome of gene expression because the novel transcripts have a different and generally lower coding potential. This was clearly demonstrated for the *RXT3* locus, where we directly showed that an overall higher RNAPII CRAC and RNAseq signals translate into lower levels of full length protein. Conversely, for *APS2* an apparently lower transcriptional signal translates into higher levels of protein, because the TSS shift favors a lower abundance species with a higher coding potential. This already high level of indetermination is further complicated by some uncertainty concerning the fate of the new transcripts that contain premature stop codons or upstream ORFs. Although we show that these RNAs are generally strongly sensitive to NMD, some of them can still be detected in wild-type cells. It is unclear why they escape NMD, but it is possible that some undergo frame-shifting during translation and produce proteins with similar or identical composition as the normal gene product.

Note that our data do not exclude a more traditional role in transcription for Rap1 and other GRFs as activators, possibly mediated by their activation domains, or as repressors. Indeed, we found cases of gene downregulation in the absence of Rap1 that were not associated with detectable nucleosome changes (Figures 3G-H, bottom region of the heatmap; data not shown) and were not suppressed by expression of Rap1_DBD_.

A similar discrepancy between RNA abundance and protein levels was the starting point of a recent study demonstrating the regulation of a meiotic network by the use of alternative TSSs producing isoforms with different coding potential (Cheng et al., 2018). Although the mechanisms by which the TSS shifts are generated is likely to be different, the general conclusions of this study are similar.

These findings have important implications for the mathematical modeling of complex gene expression networks. Such holistic approaches are often based on the assumption that RNA abundance or RNAPII occupancy directly translate into the expression of a gene product that can in turn influence the network. This is, however, not that frequently verified, and many physiological and non-physiological perturbations might bring about changes in transcription patterns similar to the ones that we describe here, which could be source of inaccuracy for gene network modeling.

## ACKNOWLEDGEMENTS

We wish to thank K. Adelman, U. Aiello, A.Oldfield, J.Gros, O. Porrua, M. Werner and T. Villa for critical reading of the manuscript, and K. Adelman, A. Oldfield, R. Jothi and F. van Werven for sharing results before publications. We also wish to thank Yan Jaszczyszyn for expert technical help in preparing CRAC libraries for sequencing. This work was supported by the Centre National de la Recherche Scientifique (C.N.R.S.), the Fondation pour la Recherche Medicale (F.R.M., programme équipes 2013), l’Agence National pour la Recherche (A.N.R., grants ANR-12-BSV8-0014-01 and ANR-16-CE12-0022-01 to D.L.), the Labex Who Am I? (ANR-11-LABX-0071 et Idex ANR-11-IDEX-0005-02 to D.L.) and the Swiss National Fund (grant no. 31003A_170153 to D.S.). D.C. and T.C. have been supported by fellowships from the French Ministry of Research. This work has benefited from the facilities and expertise of the high throughput sequencing core facility of I2BC (Centre de Recherche de Gif - http://www.i2bc-saclay.fr/).

## AUTHORS CONTRIBUTION

Conceptualization: D.L., D.C., S.K.; Methodology: T.C., M.B., D.C.; Software: M.B., T.C.; Formal Analysis: M.B., D.L., D.C.; Investigation: D.C., S.K., F.F., C.B., H.G.; Writing – Original Draft: D.C., D.L.; Writing, Review and Editing: D.C., D.L., S.K., D.S.; Funding Acquisition: D.L., D.S.; Supervision: D.L.

## METHODS

### Yeast strains, plasmids and oligonucleotides

Yeast strains, plasmids and oligonucleotides used in this study are described in Supplemental Tables 1-3.

### RNA and protein analysis

Northern and western blot analyses were performed with standard protocols. Probes used for Northern blot hybridization are listed in Supplemental Table 3

### RNAPII CRAC

RNAPII CRAC data upon nuclear depletion of Rap1 and Reb1 have been generated in a separate study (Candelli et al., 2018). For nuclear depletion of Abf1 rapamycin was added to the Abf1 anchor away strain for 30 and 90 minutes and data from the 90 minutes time point was used for the analyses shown in Figure 6. Rap1 was also depleted using the auxin degron system (Nishimura et al., 2009)(Nishimura et al., 2009) by adding IAA (Indole-3’-Acetic Acid, Sigma) 500*µ*M to Rap1-AID cells for 10 or 20min before crosslinking.

The CRAC protocol used in this study is derived from Granneman et al. (Granneman et al., 2009), modified as described in (Candelli et al., 2018). Briefly, 2L of yeast cells expressing Rpb1-HTP tag were grown at 30°C to OD_600_=0.6 in CSM-Trp medium before addition of rapamycin or IAA for the times required. Cells were UV crosslinked using a W5 UV crosslinking unit (UVO3 Ltd) for 50 seconds, harvested by centrifugation, washed in cold PBS and resuspended in TN150 buffer (50 mM Tris pH 7.8, 150 mM NaCl, 0.1% NP-40 and 5 mM beta mercaptoethanol, 2.4 ml/g of cells) supplemented with protease inhibitors (Complete(tm), EDTA-free Protease Inhibitor Cocktail). The suspension was flash frozen in droplets and cells were mechanically broken with a Mixer Mill MM 400 (5 cycles of 3 minutes at 20 Hz). Extracts were treated for one hour at 25°C with DNase I (165U/g of cells) to solubilize chromatin and then clarified by centrifugation (20min at 20000g at 4°C). The complexes were purified by a two step procedure, the second one under denaturing conditions as described (Candelli et al., 2018). High salt washes for both purification steps were done at 1 M NaCl for increased stringency. The dephosphorylation step required for cleaving the 2’-5’ cyclic phosphate left by RNase treatment (Granneman et al., 2009) was omitted to enrich for nascent transcripts, the 3’ end of which being protected from RNAse treatment. The protein fractionation step was performed with a Gel Elution Liquid Fraction Entrapment Electrophoresis (GelFree) system (Expedeon). Rpb1-containing fractions were treated with 100 *µ*g of proteinase K in a buffer containing 0.5 % SDS. RNAs were purified and reverse transcribed using reverse transcriptase Superscript IV (Invitrogen).

The concentration of cDNAs in the reaction was estimated by quantitative PCR using a standard of known concentration. PCR amplification were performed in separate 25 *µ*l reactions containing each 2 *µ*l of cDNA for typically 7-9 PCR cycles (LA Taq, Takara). The PCR reactions from all the samples were pooled and treated for 1 hour at 37°C with 200 U/ml of Exonuclease I (NEB). The DNA was purified using NucleoSpin^®^ Gel and PCR Clean-up (Macherey-Nagel) and sequenced using Illumina technology.

### TSS sequencing

The TSS sequencing protocol has been described in Malabat et al. (Malabat et al., 2015). Yeast cells were grown to mid-exponential phase in YPD-rich media. After treatment of cells with Rapamycin, Auxin, both or none, *Schizosaccharomyces pombe* cells were added for spiking purposes in a 1 to 10 ratio. Cells were then harvested and pellets were frozen in liquid nitrogen. Total RNA was extracted using two successive hot phenol steps and one chloroform. After ethanol precipitation, RNA pellet was treated with DNase and extracted again with phenol chloroform 5:1 pH4.5.

Polyadenylated transcripts were purified from 75 *µ*g of total RNAs using oligo d(T)_25_magnetic beads (New England Biolabs). RNAs were dephosphorylated using FastAP Thermosensitive Alkaline Phosphatase (ThermoFisher) and treated with Cap-Clip Acid Pyrophosphatase (Tebu-bio). RNAs were then ligated overnight at 16°C to the biotinylated 5’ adaptor (oligonucleotide 3365, 50pmol) using T4 RNA ligase I (10 units, New England Biolabs) and ATP at a final concentration of 1mM. After 5’ ligation, the RNA was fragmented for 5’ at 70°C in fragmentation buffer (10mM ZnCl_2_, 10mM Tris pH7.5). The reaction was stopped by adding 75 *µ*l of a cold solution containing 1 *µ*l of EDTA 0.5M. Ligated RNA molecules were purified on streptavidin magnetic beads from New England Biolabs (50 *µ*l of slurry). Binding was performed at 37°C for 10’, beads were washed and RNA were eluted in 20*µ*l of water at 95°C for 5’. Reverse transcription was performed with 50pmoles of primer 3018 and 300 units of RevertAid reverse transcriptase (ThermoFisher) in 30 *µ*l. cDNAs were purified by adding 1.8 volumes of Agencourt RNAClean XP beads (Beckman Coulter) and eluted in 50 *µ*l of water at room temperature for 10’.

4 independent PCR reactions were performed in a final volume of 25 *µ*l with 5 pmoles each of primer 1 and Illumina multiplexing PCR primer, and 0.25 *µ*l of LA Taq DNA polymerase (Takara). The four PCR reactions were pooled, purified on NucleoMag NGS Clean-up and Size Select dynabeads (Macherey-Nagel) and the DNA eluted in 30 *µ*l of water.

### MNase-Seq

MNase-Seq has been performed based on the procedure describe in Kubik et al. (Kubik et al., 2015a). Briefly, 120 ml of Rap1-AA yeast cells ectopically expressing Rap1_DBD_, wild-type Rap1 or containing an empty plasmid were grown at 30°C in CSM-His media to OD_600_∼0.15. Rapamycine was added for one hour to a final concentration of 1*µ*g/ml and cells were crosslinked for 5min in 1% formaldehyde at room temperature. After crosslinking the procedure was carried out as described (Kubik et al., 2015)

### Dataset processing and data analysis

CRAC datasets were analyzed as described (Candelli et al., 2018). The pyCRAC script pyFastqDuplicateRemover was used to collapse PCR duplicates using a 6 nucleotides random tag included in the 3’ adaptor (supplemental table 2). The resulting sequences were reverse complemented with Fastx reverse complement (part of the fastx toolkit, http://hannonlab.cshl.edu/fastx_toolkit/) and mapped to the R64 genome (Cherry et al., 2012) with bowtie2 (-N 1 {f) (Langmead and Salzberg, 2012).

RNA-seq and TSS-seq samples were demultiplexed by the sequencing platform with bcl2fastq2 v2.15.0 and illumina trueseq adaptors were trimmed with cutadapt 1.9. Sequencing reads were quality trimmed with trimmomatic and mapped to the R64 genome with bowtie2 (default options).

Data obtained from RNAPII CRAC, TSS Seq, RNA Seq and MNase Seq were analyzed using deeptools 2.0 (Ramírez et al., 2014) on the Roscoff (http://galaxy3.sb-roscoff.fr/) and Freiburg (http://deeptools.ie-freiburg.mpg.de/) Galaxy platforms. The list of sites used for each analysis can be found in Supplementary table 4.

All data generated in this report are available under GEO number GSE114589. RNAPII CRAC Data for Reb1-AA and Rap1-AA and RNA-Seq data for the Rap1_DBD_ series have been generated in a previous study (Candelli et al., 2018).

## REFERENCES

Azad, G.K., and Tomar, R.S. (2016). The multifunctional transcription factor Rap1: a regulator of yeast physiology. Front. Biosci. Landmark Ed. 21, 918–930.

Badis, G., Chan, E.T., van Bakel, H., Pena-Castillo, L., Tillo, D., Tsui, K., Carlson, C.D., Gossett, A.J., Hasinoff, M.J., Warren, C.L., et al. (2008). A library of yeast transcription factor motifs reveals a widespread function for Rsc3 in targeting nucleosome exclusion at promoters. Mol. Cell 32, 878–887.

Bendjennat, M., and Weil, P.A. (2008). The transcriptional repressor activator protein Rap1p is a direct regulator of TATA-binding protein. J. Biol. Chem. 283, 8699–8710.

Bosio, M.C., Fermi, B., and Dieci, G. (2017). Transcriptional control of yeast ribosome biogenesis: A multifaceted role for general regulatory factors. Transcription 8, 254–260.

Candelli, T., Challal, D., Briand, J.-B., Boulay, J., Porrua, O., Colin, J., and Libri, D. (2018). High-resolution transcription maps reveal the widespread impact of roadblock termination in yeast. EMBO J. 37.

Cheng, Z., Otto, G.M., Powers, E.N., Keskin, A., Mertins, P., Carr, S.A., Jovanovic, M., and Brar, G.A. (2018). Pervasive, Coordinated Protein-Level Changes Driven by Transcript Isoform Switching during Meiosis. Cell 172, 910–923.e16.

Chereji, R.V., Ramachandran, S., Bryson, T.D., and Henikoff, S. (2018). Precise genome-wide mapping of single nucleosomes and linkers in vivo. Genome Biol. 19, 19.

Cherry, J.M., Hong, E.L., Amundsen, C., Balakrishnan, R., Binkley, G., Chan, E.T., Christie, K.R., Costanzo, M.C., Dwight, S.S., Engel, S.R., et al. (2012). Saccharomyces Genome Database: the genomics resource of budding yeast. Nucleic Acids Res. 40, D700–705.

Fishburn, J., Galburt, E., and Hahn, S. (2016). Transcription Start Site Scanning and the Requirement for ATP during Transcription Initiation by RNA Polymerase II. J. Biol. Chem. 291, 13040–13047.

Ganapathi, M., Palumbo, M.J., Ansari, S.A., He, Q., Tsui, K., Nislow, C., and Morse, R.H. (2011). Extensive role of the general regulatory factors, Abf1 and Rap1, in determining genome-wide chromatin structure in budding yeast. Nucleic Acids Res. 39, 2032–2044.

Garbett, K.A., Tripathi, M.K., Cencki, B., Layer, J.H., and Weil, P.A. (2007). Yeast TFIID serves as a coactivator for Rap1p by direct protein-protein interaction. Mol. Cell. Biol. 27, 297–311.

Giesman, D., Best, L., and Tatchell, K. (1991). The role of RAP1 in the regulation of the MAT alpha locus. Mol. Cell. Biol. 11, 1069–1079.

Granneman, S., Kudla, G., Petfalski, E., and Tollervey, D. (2009). Identification of protein binding sites on U3 snoRNA and pre-rRNA by UV cross-linking and high-throughput analysis of cDNAs. Proc. Natl. Acad. Sci. U. S. A. 106, 9613–9618.

Hartley, P.D., and Madhani, H.D. (2009). Mechanisms that specify promoter nucleosome location and identity. Cell 137, 445–458.

Haruki, H., Nishikawa, J., and Laemmli, U.K. (2008). The anchor-away technique: rapid, conditional establishment of yeast mutant phenotypes. Mol. Cell 31, 925–932.

Hughes, A.L., Jin, Y., Rando, O.J., and Struhl, K. (2012). A functional evolutionary approach to identify determinants of nucleosome positioning: a unifying model for establishing the genome-wide pattern. Mol. Cell 48, 5–15.

Jensen, T.H., Jacquier, A., and Libri, D. (2013). Dealing with pervasive transcription. Mol. Cell 52, 473–484.

Jin, Y., Eser, U., Struhl, K., and Churchman, L.S. (2017). The Ground State and Evolution of Promoter Region Directionality. Cell 170, 889–898.e10.

Johnson, A.N., and Weil, P.A. (2017). Identification of a transcriptional activation domain in yeast repressor activator protein 1 (Rap1) using an altered DNA-binding specificity variant. J. Biol. Chem. 292, 5705–5723.

Kasahara, K., Ohyama, Y., and Kokubo, T. (2011). Hmo1 directs pre-initiation complex assembly to an appropriate site on its target gene promoters by masking a nucleosome-free region. Nucleic Acids Res. 39, 4136–4150.

Kim, J.H., Lee, B.B., Oh, Y.M., Zhu, C., Steinmetz, L.M., Lee, Y., Kim, W.K., Lee, S.B., Buratowski, S., and Kim, T. (2016). Modulation of mRNA and lncRNA expression dynamics by the Set2-Rpd3S pathway. Nat. Commun. 7, 13534.

Kim, T., Xu, Z., Clauder-Münster, S., Steinmetz, L.M., and Buratowski, S. (2012). Set3 HDAC Mediates Effects of Overlapping Noncoding Transcription on Gene Induction Kinetics. Cell 150, 1158–1169.

Knight, B., Kubik, S., Ghosh, B., Bruzzone, M.J., Geertz, M., Martin, V., Dénervaud, N., Jacquet, P., Ozkan, B., Rougemont, J., et al. (2014). Two distinct promoter architectures centered on dynamic nucleosomes control ribosomal protein gene transcription. Genes Dev. 28, 1695–1709.

Krietenstein, N., Wal, M., Watanabe, S., Park, B., Peterson, C.L., Pugh, B.F., and Korber, P. (2016). Genomic Nucleosome Organization Reconstituted with Pure Proteins. Cell 167, 709–721.e12.

Kubik, S., Bruzzone, M.J., Jacquet, P., Falcone, J.-L., Rougemont, J., and Shore, D. (2015). Nucleosome Stability Distinguishes Two Different Promoter Types at All Protein-Coding Genes in Yeast. Mol. Cell 60, 422–434.

Kubik, S., O’Duibhir, E., Jonge, W. de, Mattarocci, S., Albert, B., Falcone, J.-L., Bruzzone, M.J., Holstege, F.C.P., and Shore, D. (2018). Sequence-directed action of RSC remodeler and pioneer factors positions +1 nucleosome to facilitate transcription. BioRxiv 266072.

Kuehner, J.N., and Brow, D.A. (2006). Quantitative analysis of in vivo initiator selection by yeast RNA polymerase II supports a scanning model. J. Biol. Chem. 281, 14119–14128.

Kurtz, S., and Shore, D. (1991). RAP1 protein activates and silences transcription of mating-type genes in yeast. Genes Dev. 5, 616–628.

Lai, W.K.M., and Pugh, B.F. (2017). Understanding nucleosome dynamics and their links to gene expression and DNA replication. Nat. Rev. Mol. Cell Biol. 18, 548–562.

Langmead, B., and Salzberg, S.L. (2012). Fast gapped-read alignment with Bowtie 2. Nat. Methods 9, 357–359.

Lee, W., Tillo, D., Bray, N., Morse, R.H., Davis, R.W., Hughes, T.R., and Nislow, C. (2007). A high-resolution atlas of nucleosome occupancy in yeast. Nat. Genet. 39, 1235–1244.

Lieleg, C., Krietenstein, N., Walker, M., and Korber, P. (2015). Nucleosome positioning in yeasts: methods, maps, and mechanisms. Chromosoma 124, 131–151.

Malabat, C., Feuerbach, F., Ma, L., Saveanu, C., and Jacquier, A. (2015). Quality control of transcription start site selection by nonsense-mediated-mRNA decay. ELife 4.

Marquardt, S., Escalante-Chong, R., Pho, N., Wang, J., Churchman, L.S., Springer, M., and Buratowski, S. (2014). A chromatin-based mechanism for limiting divergent noncoding transcription. Cell 157, 1712–1723.

Murakami, K., Mattei, P.-J., Davis, R.E., Jin, H., Kaplan, C.D., and Kornberg, R.D. (2015). Uncoupling Promoter Opening from Start-Site Scanning. Mol. Cell 59, 133–138.

Neil, H., Malabat, C., d’Aubenton-Carafa, Y., Xu, Z., Steinmetz, L.M., and Jacquier, A. (2009). Widespread bidirectional promoters are the major source of cryptic transcripts in yeast. Nature 457, 1038–1042.

Nishimura, K., Fukagawa, T., Takisawa, H., Kakimoto, T., and Kanemaki, M. (2009). An auxin-based degron system for the rapid depletion of proteins in nonplant cells. Nat. Methods 6, 917–922.

Papai, G., Tripathi, M.K., Ruhlmann, C., Layer, J.H., Weil, P.A., and Schultz, P. (2010). TFIIA and the transactivator Rap1 cooperate to commit TFIID for transcription initiation. Nature 465, 956–960.

Parnell, T.J., Huff, J.T., and Cairns, B.R. (2008). RSC regulates nucleosome positioning at Pol II genes and density at Pol III genes. EMBO J. 27, 100–110.

Parnell, T.J., Schlichter, A., Wilson, B.G., and Cairns, B.R. (2015). The chromatin remodelers RSC and ISW1 display functional and chromatin-based promoter antagonism. ELife 4, e06073.

Porrua, O., and Libri, D. (2015). Transcription termination and the control of the transcriptome: why, where and how to stop. Nat Rev Mol Cell Biol.

Preti, M., Ribeyre, C., Pascali, C., Bosio, M.C., Cortelazzi, B., Rougemont, J., Guarnera, E., Naef, F., Shore, D., and Dieci, G. (2010). The telomere-binding protein Tbf1 demarcates snoRNA gene promoters in Saccharomyces cerevisiae. Mol. Cell 38, 614–620.

Ramírez, F., Dündar, F., Diehl, S., Grüning, B.A., and Manke, T. (2014). deepTools: a flexible platform for exploring deep-sequencing data. Nucleic Acids Res. 42, W187–191.

Reid, J.L., Iyer, V.R., Brown, P.O., and Struhl, K. (2000). Coordinate regulation of yeast ribosomal protein genes is associated with targeted recruitment of Esa1 histone acetylase. Mol. Cell 6, 1297–1307.

Reja, R., Vinayachandran, V., Ghosh, S., and Pugh, B.F. (2015). Molecular mechanisms of ribosomal protein gene coregulation. Genes Dev. 29, 1942–1954.

Rhee, H.S., and Pugh, B.F. (2012). Genome-wide structure and organization of eukaryotic pre-initiation complexes. Nature 483, 295–301.

Ryan, M.P., Jones, R., and Morse, R.H. (1998). SWI-SNF complex participation in transcriptional activation at a step subsequent to activator binding. Mol. Cell. Biol. 18, 1774–1782.

Shivaswamy, S., and Iyer, V.R. (2008). Stress-dependent dynamics of global chromatin remodeling in yeast: dual role for SWI/SNF in the heat shock stress response. Mol. Cell. Biol. 28, 2221–2234.

Struhl, K., and Segal, E. (2013). Determinants of nucleosome positioning. Nat. Struct. Mol. Biol. 20, 267–273.

Tomar, R.S., Zheng, S., Brunke-Reese, D., Wolcott, H.N., and Reese, J.C. (2008). Yeast Rap1 contributes to genomic integrity by activating DNA damage repair genes. EMBO J. 27, 1575–1584.

Tsankov, A.M., Thompson, D.A., Socha, A., Regev, A., and Rando, O.J. (2010). The role of nucleosome positioning in the evolution of gene regulation. Plos Biol. 8, e1000414.

Whitehouse, I., Rando, O.J., Delrow, J., and Tsukiyama, T. (2007). Chromatin remodelling at promoters suppresses antisense transcription. Nature 450, 1031–1035.

Wu, A.C., Patel, H., Chia, M., Moretto, F., Frith, D., Snijders, A.P., and Werven, F.J. van (2018). Repression of divergent noncoding transcription by a sequence-specific transcription factor. BioRxiv 314310.

Xu, Z., Wei, W., Gagneur, J., Perocchi, F., Clauder-Munster, S., Camblong, J., Guffanti, E., Stutz, F., Huber, W., and Steinmetz, L.M. (2009). Bidirectional promoters generate pervasive transcription in yeast. Nature 457, 1033–1037.

Yarragudi, A., Miyake, T., Li, R., and Morse, R.H. (2004). Comparison of ABF1 and RAP1 in chromatin opening and transactivator potentiation in the budding yeast Saccharomyces cerevisiae. Mol. Cell. Biol. 24, 9152–9164.

Yu, L., Sabet, N., Chambers, A., and Morse, R.H. (2001). The N-terminal and C-terminal domains of RAP1 are dispensable for chromatin opening and GCN4-mediated HIS4 activation in budding yeast. J. Biol. Chem. 276, 33257–33264.

